# Frequency Effects on Spelling Error Recognition: An ERP Study

**DOI:** 10.1101/2022.01.07.475328

**Authors:** Ekaterina V. Larionova, Olga V. Martynova

## Abstract

Spelling errors are ubiquitous in all writing systems. Most studies exploring spelling errors focused on the phonological plausibility of errors. However, unlike typical pseudohomophones, spelling errors occur in naturally produced written language with variable frequencies. We investigated the time course of recognition of the most frequent orthographic errors in Russian (error in an unstressed vowel at the root) and the effect of word frequency on this process. During ERP recording, 26 native Russian speakers silently read high-frequency correctly spelled words, low-frequency correctly spelled words, high-frequency words with errors, and low-frequency words with errors. The amplitude of P200 was more positive for correctly spelled words than for misspelled words and did not depend on the frequency of the words. Word frequency affected spelling recognition in the later stages of word processing (350–700 ms): high-frequency misspelled words elicited a greater P300 than high-frequency correctly spelled words, and low-frequency misspelled words elicited a greater N400 than low-frequency correctly spelled words. We observe spelling effects in the same time window for both the P300 and N400, which may reflect temporal overlap between mainly categorization processes based on orthographic properties for high-frequency words and phonological processes for low-frequency words. We concluded that two independent pathways can be active simultaneously during spelling recognition: one reflects mainly orthographic processing of high-frequency words and the other is the phonological processing of low-frequency words. Our findings suggest that these pathways are associated with different ERP components. Therefore, our results complement existing reading models and demonstrate that the neuronal underpinnings of spelling error recognition during reading depend on word frequency.

## 1 Introduction

Reading speed and efficiency are achieved through automatic visual word recognition. Reading includes the visual encoding of letters, transforming letters into graphemes and orthographic patterns, lexical and phonological analysis, and understanding the meaning of written words (Grainger and Jacobs, 1996; Bentin et al., 1999). Spelling errors are typically thought of as a footprint of a word’s weak orthographic representation in the mind, and the cause of their occurrence is a lack of stability in a word’s spelling, sound, or semantics (Perfetti, 1985; Perfetti and Hart, 2001). Most studies on spelling errors focused on the errors’ phonological plausibility (Landerl and Wimmer, 2000; Caravolas and Volín, 2001; Quémart and Casalis, 2017). Misspelled words are indeed pseudohomophones: they differ from words in their orthography but have the same phonology, for example, pseudohomophone SPAIS and word SPACE in English. The pseudohomophone effect is a well-studied marker of phonological activation and comprises a higher error rate and response time for pseudohomophones compared to base words; in addition, pseudohomophones are more often falsely classified as words, presumably due to correct phonological representation (Rubenstein et al., 1971; van Orden, 1987; Goswami et al., 2001; Pexman et al., 2001; Briesemeister et al., 2009). Rahmanian and Kuperman (2019) noted that spelling errors are fundamentally different from typical pseudohomophones: they occur in naturally produced written language with variable frequencies. However, the use of basic spelling errors as pseudohomophone stimuli is relatively rare.

The frequency effect is a well-studied marker of lexical access. Word frequency is one of the strongest predictors of word processing efficiency (Monsell et al., 1989; Brysbaert et al., 2018). High-frequency words are familiar to more people and are processed faster than low-frequency words (Monsell et al., 1989). The frequency effect has been identified in many languages: English (Rayner, 1998), Portuguese (Faísca et al., 2019), French (Segui et al., 1982), Chinese (Yan et al., 2006), and Russian (Laurinavichyute et al., 2019). Word frequency influences the temporal course of semantic and phonological activation in word recognition (Zhang et al., 2009). High-frequency words are more susceptible to phonetic changes (Bondarko et al., 1988; Raeva and Riekhakaynen, 2016; Todd et al., 2019); in addition, there is a hypothesis that the uncertainty created by competition between different spellings will be greater for high-frequency words (Rahmanian and Kuperman, 2019). Therefore, error recognition in words of different frequencies may differ, and thus, it is possible that error recognition in high-frequency words may be even more difficult than in low-frequency words.

Concerning the frequency effect in visual word recognition, the ERP evidence is consistent. Generally, low-frequency words elicited larger ERP amplitudes than high-frequency words (Osterhout et al., 1997; Sereno et al., 1998, 2003; Assadollahi and Pulvermüller, 2003; Hauk and Pulvermüller, 2004; Hauk et al., 2006). The frequency effect was found both in the presentation of words in isolation (Sereno et al., 1998) and in the context of sentences (Sereno et al., 2003). Larger amplitudes for low-frequency words reflect the difficulty in accessing their lexical representations (Sereno et al., 2003). Using MEG, Assadollahi and Pulvermuller (2003) showed the dependence of the frequency effect on word length during reading recurrent words: the frequency effect was observed 120–170 ms after stimulus onset for short words of 3–4 letters and after 225–250 ms for long words of 5–7 letters. The localization of the source showed that the effect of frequency was most pronounced over the left occipitotemporal areas (visual word form areas) (Assadollahi and Pulvermüller, 2003). Osterhout et al. (1997) reported a frequency effect ranging from 150 to 250 ms during reading normal or scrambled English prose; the average length of their stimuli ranged from 2.5 to 6.2 letters. Hauk et al. (2006) showed an early frequency effect at 110 ms (for words of 3–6 letters) using a visual lexical decision task. In the experiment of Proverbio et al. (2008), the frequency effect was observed later, at around 240 ms, which the authors associated with the presence of long words (8–9 letters) in the stimulus set. Furthermore, low-frequency words showed lower amplitudes than high-frequency ones, which does not align with the results of other studies; the reason for this result, according to the authors, was the specific nature of the letter detection task (since no semantic analysis was required for the task) they used (Proverbio et al., 2008). Strijkers et al. (2015) investigated how different paradigms affect the frequency effect: in the semantic task, the frequency effect was observed as early as 120 ms after stimulus onset, but when categorizing the colored font of the very same words in the color task, the frequency effect emerged at the earliest 200 ms after stimulus onset.

Despite the consistency of amplitude effects of word frequency, the timing of the word frequency effect is quite discrepant across the literature, ranging from an early N1-P2 sensitivity (Sereno et al., 1998, 2003; Assadollahi and Pulvermüller, 2003; Dambacher et al., 2006; Hauk et al., 2006; Zhang et al., 2009; Araújo et al., 2015; Eberhard-Moscicka et al., 2016; Wang et al., 2021) to a longest of 300 ms and more (Polich and Donchin, 1988; Hauk and Pulvermüller, 2004; Faísca et al., 2019) even in similar experimental paradigms, indicating that lexical access is located later on in processing. For example, Hauk and Pulvermüller (2004) showed that lower ERP amplitudes in English native speakers during lexical decision tasks were elicited by words with high frequency compared to low-frequency words in the latency ranges 150–190 ms and also in 320–360 ms. The frequency effect has often been associated with N400, a negative wave between 300 and 600 ms after the stimulus (Kutas and Federmeier, 2000, 2011). The N400 amplitude is larger for low-frequency words than high-frequency words (Barber et al., 2004; Kutas and Federmeier, 2011; Vergara-Martínez et al., 2020).

The pseudohomophone effect is commonly explained by an orthography–phonology conflict (Ziegler et al., 2001; Briesemeister et al., 2009). The pseudohomophone effect was studied using ERP much less often than the frequency effect. This effect is primarily associated with phonological processing, which is found in the N400 component or even later in most studies when smaller negativity is related to higher activation of a particular phonological representation (Kramer and Donchin, 1987; Bentin et al., 1999; Proverbio et al., 2004; Vissers et al., 2006; Briesemeister et al., 2009; González-Garrido et al., 2015; Costello et al., 2021). N400 has a larger amplitude for pseudohomophones than for words (Briesemeister et al., 2009; Hasko et al., 2013; González-Garrido et al., 2015). However, the ERP results show that phonological activation may occur at an early stage of visual word recognition as early as 150 ms (associated with the P2 component) after stimulus onset and may influence lexical access (Braun et al., 2009; Zhang et al., 2009). Unlike the frequency effect, the pseudohomophone effect does not depend on the length of the stimulus (Ziegler et al., 2001; Briesemeister et al., 2009). However, phonological effects can be modulated by the orthographic transparency of the writing system (Simon et al., 2006; Frost and Ziegler, 2007).

Some previous ERP studies using pseudohomophone stimuli are summarized in Table 1. We would like to draw attention to the conditions for generating pseudohomophone stimuli, which were not always described in detail, and the frequency of the basic words used to create pseudohomophones. To generate pseudohomophones, both one and two letters of the base word were changed (e.g., Braun et al., 2009; Briesemeister et al., 2009; Costello et al., 2021); these were only vowels (e.g., Vissers et al., 2006) or vowels and consonants at the same time (e.g., Briesemeister et al., 2009). Only González-Garrido et al. (2015) mentioned that the incentives used are the most frequent orthographic errors in Spanish. Some authors pay attention to the fact that their pseudohomophones are not orthographically similar to words (e.g., Newman and Connolly, 2004) to minimize the contribution of orthography to their processing. Using words with fundamental spelling errors as pseudohomophones is likely to complicate recognition, as we often face misspelled words in our daily lives instead of artificially generated pseudohomophones. The frequency of pseudohomophones was taken into account only in one experiment (Braun et al., 2009): the pseudohomophone effect was strongest for stimuli derived from low-frequency base words, a finding consistent with some previous behavioral research (Jared and Seidenberg, 1991). Therefore, we assume that frequency can influence the process of recognizing fundamental spelling errors.

**Table 1.** Summary of previous ERP studies using pseudohomophone stimuli

| **Author, year** | **Language** | **Pseudohomophones Stimuli (PsHs)** | **Task** |
| --- | --- | --- | --- |
| Newman and Connolly, 2004 | English | The conditions for generating PsHs are not described in detail (e.g., The ship disappeared into the thick phog [fog]). | Reading sentences |
| Sauseng et al., 2004 | German | The words were chosen in such a way that a homophonic letter-altered version was possible. Examples are Physik (physics) changed to Fysik. The mean frequency of the words was 40.8 per million. | Silently reading |
| Grainger et al., 2006 | English | The conditions for generating PsHs are not described in detail. PsHs were selected based on a preliminary assessment of phonological similarity with the base word (e.g., brane-BRAIN). | Masked priming |
| Vissers et al., 2006 | Dutch | PsHs were created by changing the vowel of the second syllable, keeping phonology the same. | Reading sentences |
| Braun et al., 2009 | German | PsHs were constructed by changing only one letter at the same position. For example, the PsH SAHL was derived from the base word SAAL (room). Half of the PsHs were derived from high-frequency base words, the other half of the PsHs were derived from base words of low frequency. | Lexical decision |
| Briesemeister et al., 2009 | German | PsHs were constructed by changing (i.e., replacing, adding, or removing) one or two vowels or consonants from low-frequency base words. | Lexical decision task |
| Taha and Khateb, 2013 | Arabic | PsHs were constructed by replacing a letter in the beginning syllable or by replacing a letter in the last syllable of a real word while keeping the phonology of the word identical. | Orthographic decision task |
| Araújo et al., 2015 | Portuguese | The conditions for generating PsHs are not described in detail. | Implicit reading |
| González-Garrido et al., 2015 | Spanish | PsHs were constructed by changing only one letter: e.g., omitting the silent H, or substituting homophonic letters, such as B-V, C-S-Z, LL-Y, and G-J, but keeping the phonology of the real word (the most frequent orthographic error in Spanish). Word frequency was between 50 and 100 per million. | Lexical decision |
| Kemény et al., 2018 | German | PsHs were generated by exchanging a letter or letter-cluster by a phonologically identical letter or letter-cluster (e.g., V is substituted by F, VATER is substituted by FATER). | Silently reading |
| Costello et al., 2021 | Spanish | PsHs were constructed by replacing one letter of the target word by another letter that corresponded to the same phoneme (e.g., the pseudohomophone nobio created from the base word novio [boyfriend]), (mean log word frequency 3.96). | Semantic categorization, lexical decision |

In this study, we used fundamental spelling errors in the Russian language as incentives. We constructed all the stimuli by changing only one letter in a similar position in the word; in addition, we used only one type of spelling violation—we were interested in errors in an unstressed vowel. Vowel reduction, that is, a process that neutralizes phonological contrasts between vowels in unstressed syllables, is an essential linguistic phenomenon since vowels are the main syllabic element. Vowel reduction is one of the most characteristic features of stress-timed languages: in English, many vowels in unaccented syllables are reduced to schwa, whereas in Russian, the process appears to be more complex (Jaworski, 2010). It is worth noting that the reduction of unstressed vowels is not displayed in Russian orthography. There are five vowel phonemes in Standard Russian [i, e, a, ɔ, u]. In unstressed syllables, the five-element set is reduced to two subsystems, consisting of three elements each [i, a, u] and [i, ə, u], depending on the position in the word (Kniazev and Pozaritskaya, 2011). The Russian vowels /a/ and /o/ have the same unstressed allophones, and /e/ reduces to [i] in unstressed syllables. The vowel /u/ may also be centralized, but it does not typically merge with any other vowel. We selected words with vowel phonemes [i, e, a, ɔ] in unstressed syllables, which are more susceptible to confusion. First, this choice was driven by the fact that this is a fundamental type of spelling error for the Russian language: it is pretty complex and widespread. Mistakes in unstressed vowels / i, e, o, a / are the most common, even among children with a high level of spelling competence and among foreigners studying Russian (Padgett and Tabain, 2005; Oglezneva et al., 2016; Ogneva, 2018). Second, different types of spelling violations can be recognized in different ways; for example, no one has compared the perception of pseudohomophones, which were constructed in different ways. Therefore, we have chosen only one type of error. Third, this type of error is typical not only for the Russian language but also for other East Slavic languages, implying that it is quite common throughout the world.

The lexical decision task is used most often in research on pseudohomophones (Table 1). In contrast to these studies, we used a method of silent reading, in which participants are not required to pronounce words or explicitly decide upon their lexical status. Such a paradigm allows the exploration of cognitive processes underlying reading without extraneous task demands, and according to some authors, it is better suited for research on visual word recognition (Bermúdez-Margaretto et al., 2020a; Blythe et al., 2020). We manipulated the word form frequency (high vs. low) and the correct spelling (correct words vs. words with error) of the written words in the silent word reading ERP task to test the following hypotheses: (1) word frequency influences error recognition; and (2) the time course of real error recognition is similar to pseudohomophone recognition, but due to the orthographic similarity with correctly written words, it can have its own characteristics. In this study, we aimed to investigate two ERP components, P200 and N400, which are involved in orthographic, phonological, and semantic processing and are often considered together in reading research (Dambacher et al., 2006; Zhang et al., 2009; Bermúdez-Margaretto et al., 2020b; Wang et al., 2021). To obtain behavioral data, we used a paradigm similar to the lexical decision task. We expected a faster reaction time and lower error rate for correctly written words than for misspelled words at the behavioral level.

## 2 Methods

### 2.1 Participants

Twenty-six native Russian speakers (8 males, 18 females) aged 18 to 39 years old (mean age 24.2, SD 5.3 years) with normal or corrected-to-normal vision participated in the study. All participants were students who had at least 11 years of education or had a university degree (mean 13.7, SD 2.1 years of education). They were all right-handed and did not have any reported neurological disorders or reading and spelling problems. All participants gave written informed consent in accordance with the Declaration of Helsinki; the informed consent form was approved by the Ethics Committee of the Institute of Higher Nervous Activity and Neurophysiology of the Russian Academy of Sciences (IHNA & NPh RAS).

### 2.2 Behavioral Task

#### 2.2.1 Experimental Procedure

All participants completed two tasks in a shielded room: a behavioral task and an ERP task. All stimuli were written in white Liberation Sans font 125 pt and were presented in lowercase in the center of the screen on a black background with a viewing distance of approximately 1 m. Stimuli were presented in random order on a 19” LG FLATRON L1952T monitor using the PsychoPy Experiment Builder v3.0.7 software (Peirce et al., 2019).

To evaluate the behavioral data, we asked the subjects to perform a spelling decision task with similar but not equal to ERP task stimuli (after ERP task). In contrast to the ERP task, the subjects had to give an answer regarding the correct spelling of words. Depending on the stimulus, it was necessary to press the left or right buttons of the Logitech F310 gamepad. The button used for each type of response was counterbalanced across the subjects. For 17 subjects, the left button corresponded to the correctly spelled word, and the right button corresponded to the incorrectly spelled word. For 9 subjects, the buttons were interchanged.

A stimulus was presented until the subjects’ response followed by an average interstimulus interval of 1800 ms (jittered between 1300 and 2300 ms). All of the stimuli required a response.

#### 2.2.2 Stimuli Material and Analysis

Four types of stimuli were presented: high-frequency words spelled correctly (25 words, HC—high-frequency correctly spelled words), low-frequency words spelled correctly (23 words, LC—low-frequency correctly spelled words), high-frequency words misspelled (25 words, HE—high-frequency words with errors), and low-frequency words misspelled (23 words, LE—low-frequency words with errors). All stimuli were nouns of 5–6 letters long, spelled correctly, and with errors in an unstressed vowel. The error was in the first syllable, except for two words, in which the error was in the second syllable. Example stimuli for each condition are given in Table 2. All stimuli used in this study are shown in the Appendix.

**Table 2.**
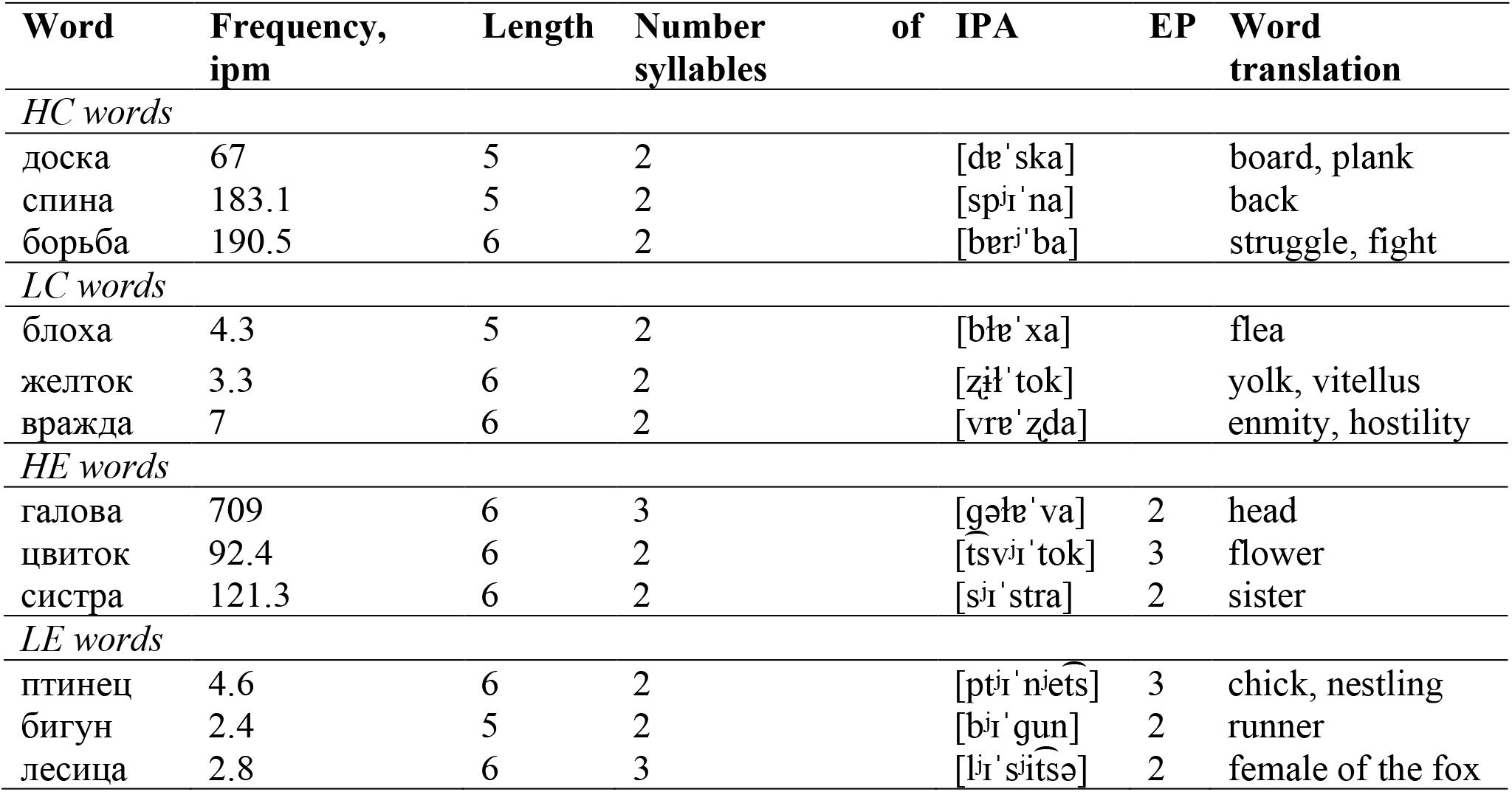
Examples of stimuli for each condition in the behavioral task. Note: ipm—instances per million words; IPA—The International Phonetic Alphabet; EP—error position for HE and LE words.

The mean frequency of high-frequency correctly spelled words was 207.77 instances per million, 152.70 instances per million for high-frequency words with errors, 12.23 instances per million for low-frequency correctly spelled words, and 8.61 instances per million for low-frequency words with errors. We used the Wilcoxon signed ranks test to confirm that there was no evidence for a reliable difference with respect to word frequency for correctly spelled words and words with errors (p > 0.1). HC and HE stimuli differed from LC and LE stimuli in word frequency of occurrence (HC vs. LC, p = 0.00003; HE vs. LE, p = 0.00003).

The reaction time and error rate for each condition were analyzed using ANOVAs with repeated measures (RM). The reaction time was evaluated regardless of the correctness of the answer. The factors were Spelling (correct vs. misspelled) and Frequency (high vs. low). In the behavioral task, the stimuli were simpler (the length was shorter and the frequency was less different between the groups of stimuli) than in the ERP task. We expected behavioral effects to be more pronounced than for ERP effects.

### 2.3 ERP Task

#### 2.3.1 Experimental Procedure

The conditions for presenting stimuli were the same as those described in 2.2.1.

During the EEG recording, the subjects had to silently read the words presented on the screen. A stimulus was shown for 200 ms, followed by an average interstimulus interval of 1850 ms (jittered between 1500 and 2200 ms). All of the stimuli required no response. The 153 stimuli were presented in two blocks, with a short break between blocks.

#### 2.3.2 Stimuli Material

All stimuli were nouns of 5–7 letters long (the mean length of each type of stimuli was 6.3 letters), spelled correctly, and with errors in an unstressed vowel, only one error was allowed in each misspelled word. There were four types of stimuli: 37 HC words, 38 LC words, 39 HE words, and 39 LE words. The error was in the first or second syllable (11 HE words and 8 LE words). Example stimuli for each condition are given in Table 3.

**Table 3.**
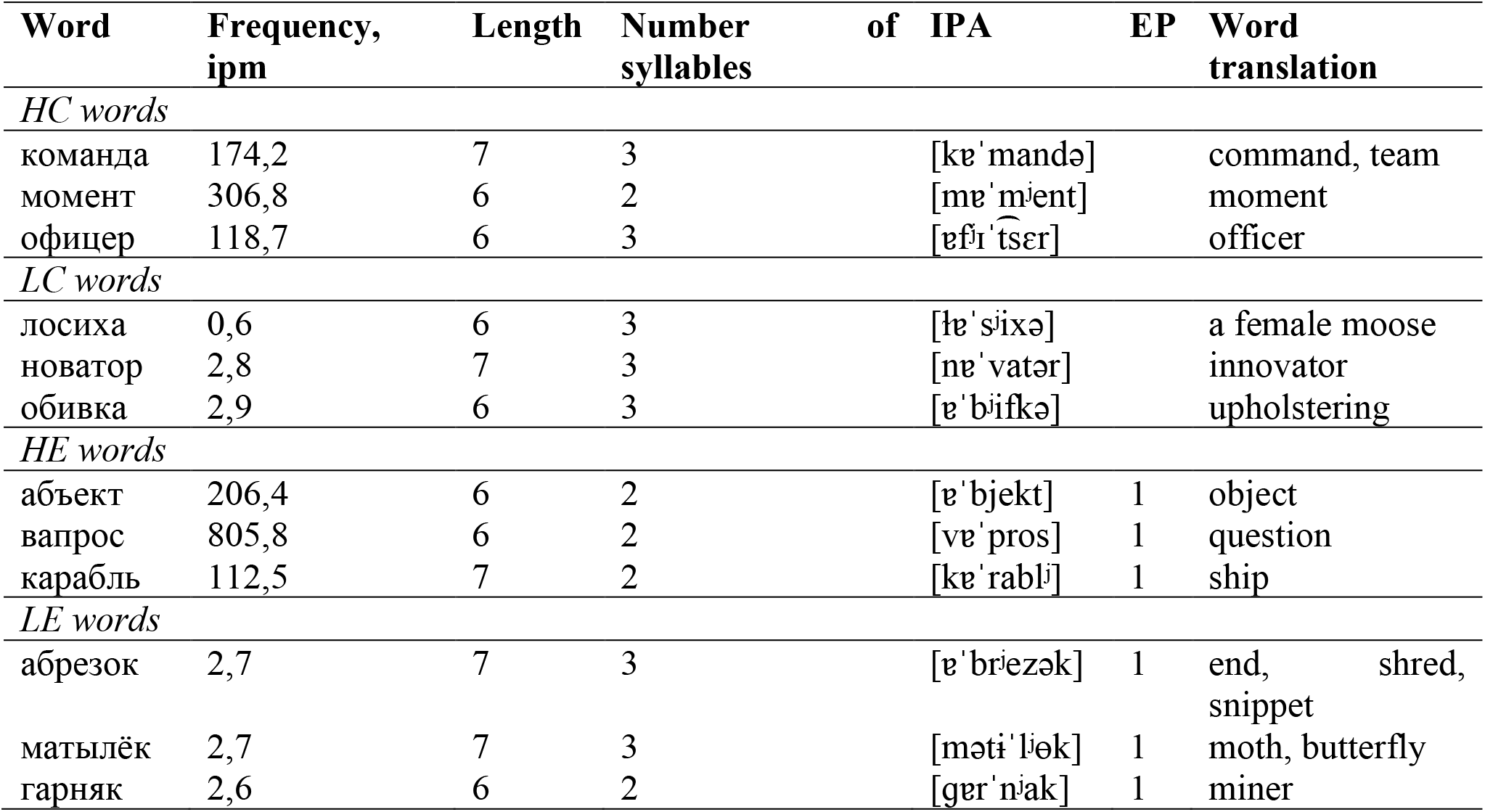
Examples of stimuli for each condition in the ERP task. Note: ipm—instances per million words; IPA—The International Phonetic Alphabet; EP—error position for HE and LE words.

All stimuli in this experiment were selected from the Frequency Dictionary of the Modern Russian Language (Lyashevskaya and Sharov, 2009). We took into account the fact that not all low-frequency words are equally difficult; for example, low-frequency compound words can consist of two high-frequency words (Brysbaert et al., 2018). All low-frequency words in the ERP and behavioral tasks were not related to high-frequency words through compounding. Since the knowledge of a word by the subject may also be an important factor in addition to the frequency of the word—the so-called variable of word prevalence introduced by Brysbaert et al. (2018)—at the end of the tasks, we asked the subjects whether they had encountered unfamiliar words. All the words in the two tasks were familiar to all subjects.

Every low-frequency word in the ERP task appears < 3 times per million, and every high-frequency word appears > 100 times per million. The mean frequency of high-frequency correctly spelled words was 258.62 instances per million, 228.38 instances per million for high-frequency words with errors, 2.45 instances per million for low-frequency correctly spelled words, and 2.70 instances per million for low-frequency words with errors. We used the Wilcoxon signed ranks test to confirm that there was no evidence for a reliable difference with respect to word frequency for correctly spelled words and words with errors (p > 0.1). HC and HE stimuli differed from LC and LE stimuli in word frequency of occurrence (HC vs. LC, p < 0.0001; HE vs. LE, p < 0.0001).

#### 2.3.3 EEG Recording and Analysis

An electroencephalogram was recorded from 19 electrodes Fp1, Fp2, F3, F4, F7, F8, C3, C4, T3, T4, T5, T6, P3, P4, O1, O2, Fz, Cz, and Pz, placed according to the International System 10-20, and reference electrodes were placed on the mastoids. Data were sampled at 250 Hz with 0.1–70 Hz filter settings with impedances below 10 kΩ. The offline processing was carried out using Brain Vision Analyzer 2.0.4 (Brain Products, GmbH, Munich, Germany). Eye movements were corrected using an ICA procedure. The data were digitally bandpass filtered (0.3–30 Hz), segmented (−300 to 1500 ms), artifact rejected (± 100 uV), followed by a visual inspection, and averaged for all of the stimuli in each condition separately. About 5% of the trials were discarded. The averaged data were baseline corrected (300 ms prior to stimulus presentation). ERPs resulted from averaging the segmented trials separately in each condition. There were 34–37 trials included for each condition in the average ERP data from most of the participants.

The analysis was performed in two independent time windows: the 160–280 ms time window (roughly corresponding to P200) and the 350–700 ms time window (roughly corresponding to N400). RM ANOVAs were applied to the average amplitude of each time window. Scalp electrodes were divided into 9 regions of interest (ROI) (Figure 1): left anterior/LA (Fp1, F7, F3), midline anterior/MA (Fz), right anterior/RA (Fp2, F4, F8), left central/LC (T3, C3), midline central/MC (Cz), right central/RC (C4, T4), left posterior/LP (T5, P3, O1), midline posterior/MP (Pz), and right posterior/RP (P4, T6, O2). We averaged the mean ERP amplitude for each ROI over the electrodes in each region. Statistical analysis was performed using the STATISTICA software (Statsoft, Tulsa, OK, USA). The factors were Spelling (correct vs. misspelled), Frequency (high vs. low), Laterality (left vs. midline vs. right), and Anterior-Posterior electrode position (anterior vs. central vs. posterior). We focused on Spelling and Frequency effects and the interactions between these factors. All significant (p < 0.05) main and interaction effects were followed by post hoc Bonferroni-corrected contrasts. To correct violations of sphericity and homogeneity, the Greenhouse–Geisser correction was applied as well. Partial eta squared (η^2^_p_) was applied as a measure of effect size, with values of 0.01–0.05 indicating small effects, 0.06–0.13 indicating medium effects, and ≥ 0.14 indicating large effects.

**Figure 1.**
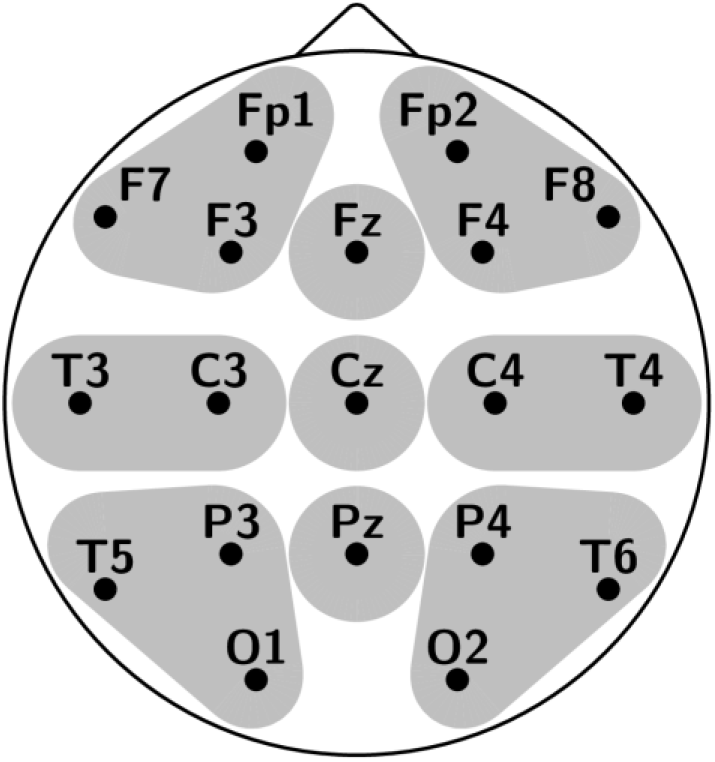
Electrode montage with regions used for analysis.

## 3 Results

### 3.1 Behavioral Data

The RM ANOVA results are shown in Figure 2. The analysis of reaction time indicated a significant main effect both for Spelling F(1,25) = 16.01, p = 0.0005, η^2^_p_ = 0.39 (correct vs. misspelled, 1.09 vs. 1.26 s) and Frequency F(1,25) = 20.48, p = 0.0001, η^2^_p_ = 0.45 (high vs. low, 1.09 vs. 1.26 s): the reaction time of low-frequency words was longer than that of high-frequency words, and the reaction time of words with errors was longer than that of correctly spelled words. No significant Spelling × Frequency interaction was found.

**Figure 2.**
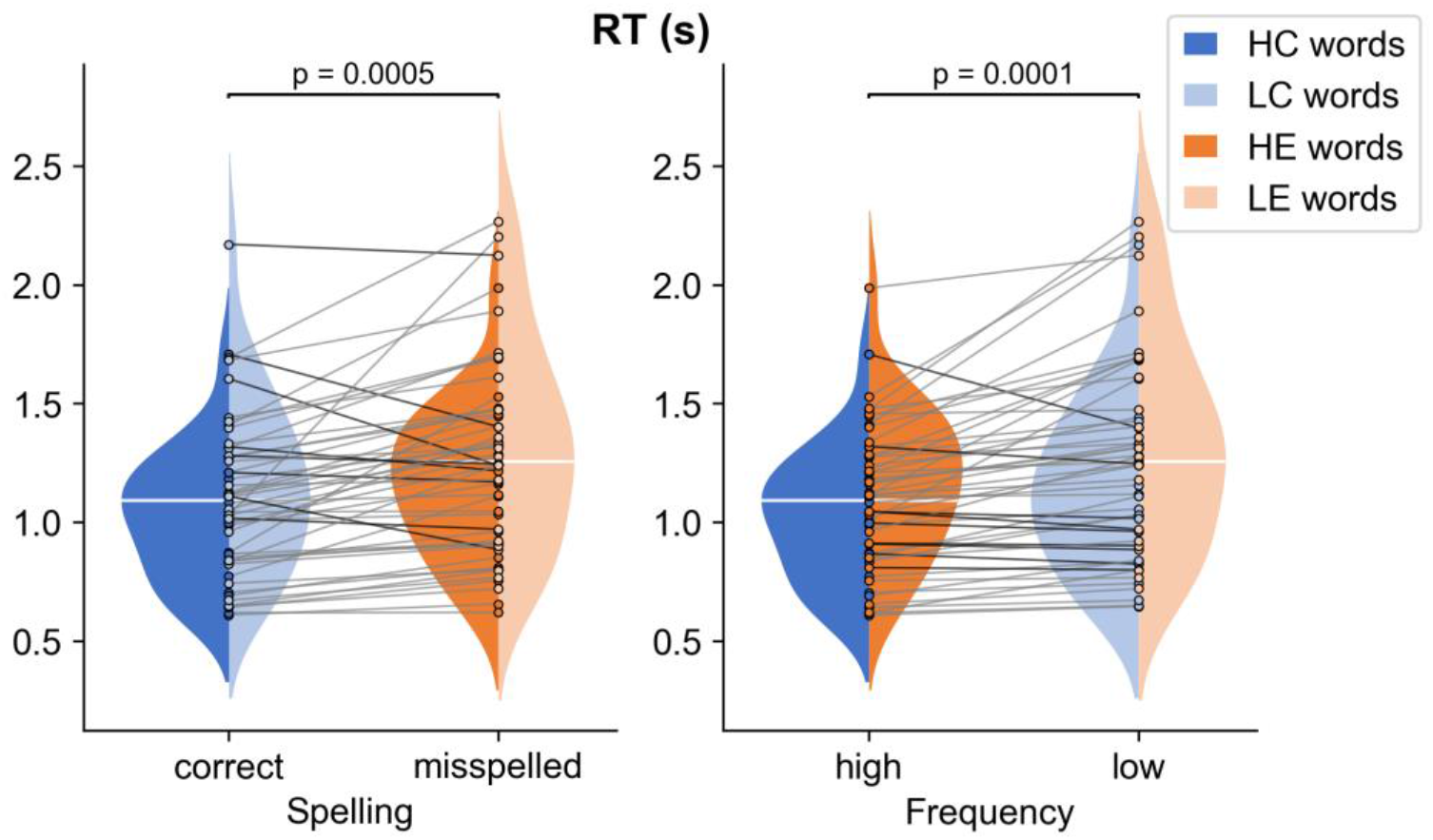
RM ANOVA results for response times (RT). The distributions of the RT for the four types of stimuli are displayed by a violin plot; means are shown as horizontal white lines.

The error rate of each condition was less than 6%: mean error rate for HC words was 0.77%, for LC words was 1.85%, for HE words was 2.62%, for LE words was 5.29%. Out of 26 participants, 21 achieved 100% accuracy for HC words and 16 achieved 100% accuracy for LC words. No further analysis was performed due to the very low number of errors.

### 3.2 ERP Data

Two time windows were selected based on data from previous studies (e.g., Bermúdez-Margaretto et al., 2020a; Wang et al., 2021 for P200; Kutas and Federmeier, 2000, 2011; Szewczyk and Schriefers, 2018, for N400) and of Global Field Power (GFP) (see Figure 3), which permits the optimal choice periods of stable topography, i.e., occurrence times of evoked components (Lehmann and Skrandies, 1980). We first computed an average ERP across each condition and participant and subjected this to GFP transformation. We found the main peaks around 160–280 ms and 350–700 ms.

**Figure 3.**
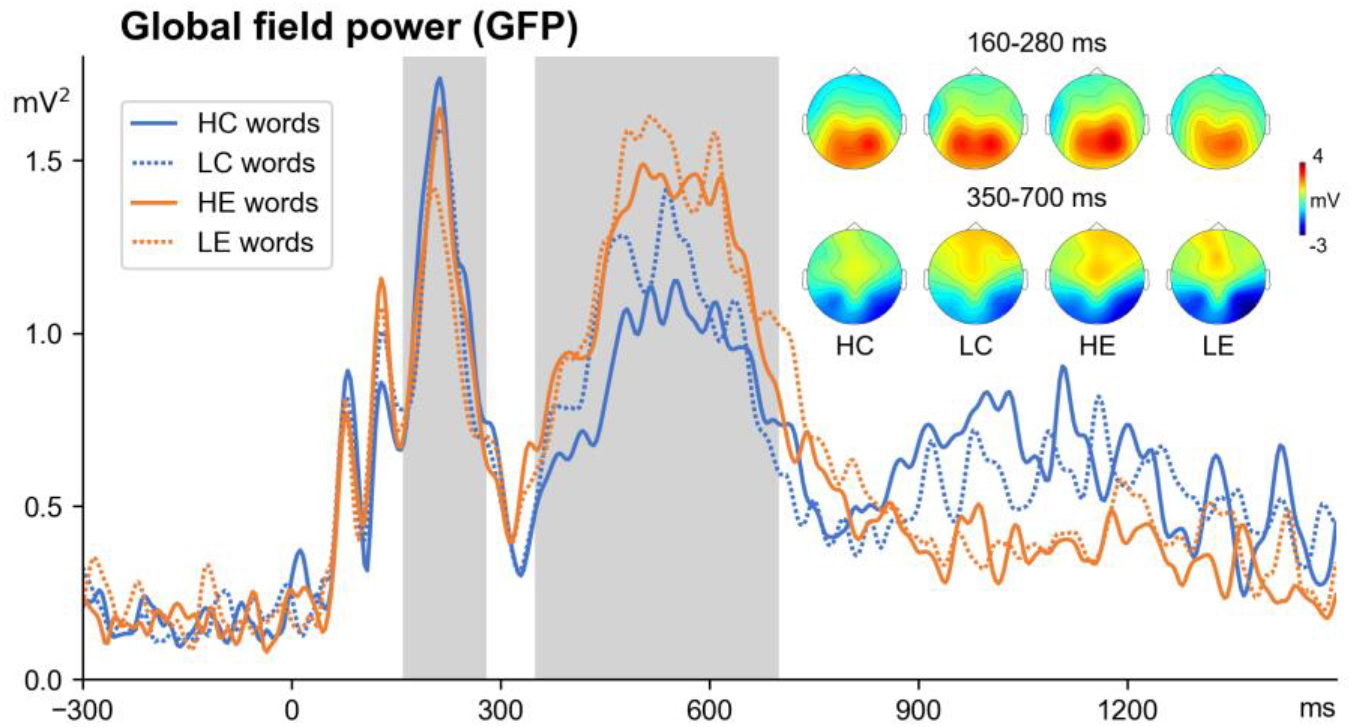
Global field power (GFP, all electrodes) averaged across the four experimental conditions and topographic maps.

Figure 4 illustrates the averaged ERPs for four conditions across all 9 regions of interest. As shown in Figure 4, the positive peak was in 160–280 ms (P200) window, and the positive and negative peaks were in the 350–700 ms (P300/N400) time windows. The timing and distribution of these components between all conditions were similar (Figure 3).

**Figure 4.**
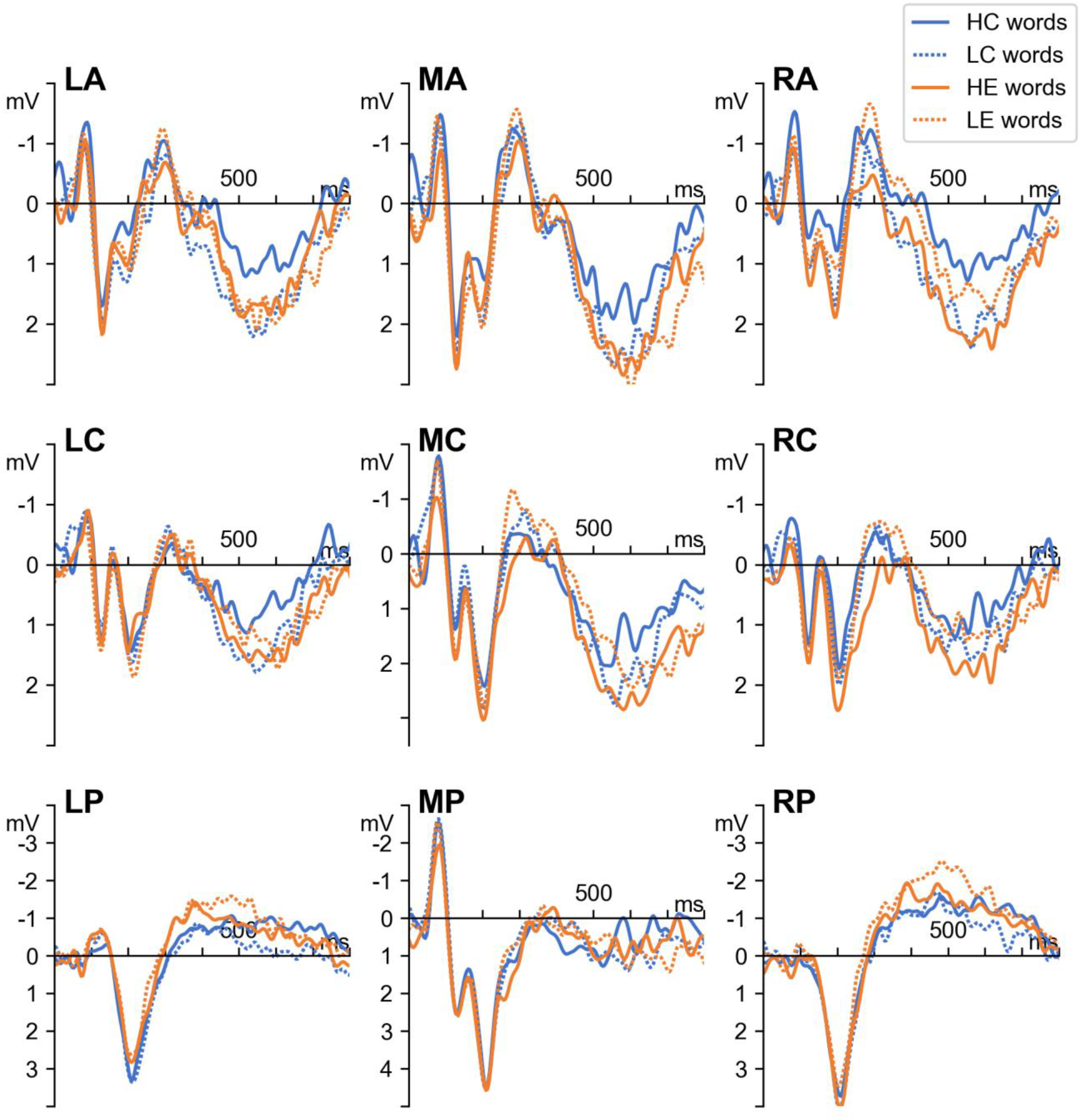
Grand average ERPs for four conditions across all 9 regions of interest.

In the 160–280-ms time window, a significant Spelling × Anteriority × Laterality interaction (F(4,100) = 3.31, p = 0.03, η^2^_p_ = 0.12) was found. Post-hoc analysis showed that the effect of Spelling was significant in the left posterior region (p = 0.0003): the amplitude of P200 was more positive for correctly spelled words than for misspelled words (2.03 vs. 1.59 μV).

In the 350–700-ms time window, a significant Spelling × Frequency interaction (F(1,25) = 4.32, p = 0.04, η^2^_p_ = 0.15) and a significant Spelling × Frequency × Anteriority × Laterality interaction (F(4,100) = 2.54, p = 0.04, η^2^_p_ = 0.10) were found. Post-hoc analysis revealed that a more positive response was for misspelled words than for correctly spelled words only for high-frequency words in the RA region (p < 0.0001, HC vs. HE, 0.71 vs. 1.75 μV) and that more negative response was for misspelled words than for correctly spelled words only for low-frequency words in the LP region (p < 0.01, LC vs. LE, −0.31 vs. −1.04 μV), RP region (p < 0.001, LC vs. LE, −0.98 vs. −1.83 μV). In addition, low-frequency correctly spelled words elicited a more positive effect than high-frequency correctly spelled words in the LA region (p < 0. 01, HC vs. LC, 0.71 vs. 1.54 μV) and RA region (p < 0.0001, HC vs. LC, 0.71 vs. 1.65 μV).

## 4 Discussion

This study demonstrates that the neuronal underpinnings of recognizing errors in words during reading depend on word frequency. High-frequency misspelled words elicited a greater P300 than high-frequency correctly spelled words, and low-frequency misspelled words elicited a greater N400 than low-frequency correctly spelled words. Meanwhile, the amplitude of P200 was more positive for correctly spelled words than for misspelled words and did not depend on the frequency of the words.

Behavioral data showed both the frequency effect (faster RT for high-frequency words than for low-frequency words) and the spelling effect (faster RT for correctly spelled words than for misspelled words). Both of these findings are consistent with the well-known word recognition effects (e.g., Monsell et al., 1989; Bentin and Ibrahim, 1996; Rayner, 1998; Proverbio et al., 2004; Yan et al., 2006; Faísca et al., 2019; Laurinavichyute et al., 2019). The lack of interaction between these factors indicates their independence from each other: error recognition speed does not depend on word frequency. Slow misspelled word processing can result from impaired grapheme-phoneme integration, since the phonological representations of such words do not coincide with their spelling representations in memory. Although we did not statistically analyze the error rate, it was highest for low-frequency misspelled words, which indicates the complexity of these stimuli for subjects. It is probably related to more blurry representations in memory for lower frequency words than for higher frequency words in which the subjects were almost not mistaken.

We found that correctly spelled words induce a larger P200 in the left posterior region than misspelled words. In the literature, P200 has been associated with word-form encoding processes and the extraction of the orthographic and phonological features of a word in the early stages of word processing (Barnea and Breznitz, 1998; Carreiras et al., 2005; Dambacher et al., 2006; Wang et al., 2021). Bles et al. (2007) suggested that P200 reflects the level of inhibition of words that mismatch phonological/orthographic input when stimuli are visually presented letter by letter. Modulation of the P200 component was found for known versus novel written words, and an increase in the amplitude of the P200 component when learning novel written words has been associated with a modification of the sublexical orthographic process and switching from letter-by-letter decoding to a more holistic lexical-type access of newly formed representations (Bermúdez-Margaretto et al., 2020b). Note that misspelled words in unstressed vowels sound like actual words, and their phonological representations do not match their orthographic representations in memory. Thus, the P200 spelling effect in our experiment can reflect the availability of orthographic representations for correctly spelled words.

The P200 distribution was broad in our study, and in contrast to the work of Bermúdez-Margaretto et al. (2020a) with a similar paradigm of passive reading, the P200 effect was observed not in the frontal-central regions but in the left temporo-parieto-occipital area. However, the topographic distribution of P200 in our study was similar to the P150 component in Gonzalez-Garrido et al.’s (2014) study with the task of categorization based on spelling, and the authors suggested that this component may index visual categorization processes sensitive to orthographic regularity. On the one hand, this difference in P200 topography may be due to the shorter presentation time of words in our study (200 ms) compared to the work of Bermúdez-Margaretto et al. (2020a). On the other hand, the neural generators of P200 are disputed, as previous studies have conflicting results, and brain source estimation revealed that the left temporal gyrus, along with the left inferior frontal gyrus, are the likeliest neural generators for P200 (Hsu et al., 2014; Bermúdez-Margaretto et al., 2020a). Furthermore, the left temporoparietal and right frontotemporal regions are the likeliest areas associated with the interaction of orthographic and phonological information in the early stages of word processing (Braun et al., 2009). FMRI data also showed that the left inferior parietal cortex supports the integration of orthography and phonology in visual word recognition (Pugh et al., 2000; Tan et al., 2005; Pattamadilok et al., 2017).

The ERP results in this experiment showed that word frequency influences the error recognition process in later stages. The N400 component found in lexical or semantic categorization tasks is overlapped by the modulation of P300, a component associated with attention mechanisms activated to perform a task (Polich, 1985, 2004; van Hees et al., 2017; Alday and Kretzschmar, 2019; Bermúdez-Margaretto et al., 2019). In our study, we also observed a positive wave in the frontal regions, along with a negative wave in the central parietal areas.

We found a more positive response for misspelled words than for correctly spelled words only for high-frequency words in the right anterior region. The P300 wave is not a component exclusively related to language processing; however, it is found in any psycholinguistic paradigm that requires an assessment of stimulus and a binary decision (Proverbio et al., 2009; González-Garrido et al., 2014; Alday and Kretzschmar, 2019). This wave reflects an information-processing cascade when attentional and memory mechanisms are engaged, namely, final stimulus evaluation (Polich, 1985, 2004; Taylor, 1993). This component can also reflect lexical decisions based on orthographic properties (Mariol et al., 2008; Kriukova and Mani, 2016). Although our passive reading task did not require making any decisions, it may involve involuntary attention processes and categorization of correctly spelled words and words with errors. Our behavioral data demonstrated that words with errors are difficult for subjects, so these stimuli impose greater demands on attention resources, which result in larger P300 amplitude. This finding is consistent with studies on stimuli with orthographic violations (e.g., Newman and Connolly, 2004; Mariol et al., 2008; González-Garrido et al., 2014) in which the P300 amplitude was higher for words or pseudowords with violations; however, these studies did not take into account word frequency. Mariol et al. (2008) reported a larger P300 around 300–550 msec on frontal and central sites for pseudowords with double consonants never doubled than pseudowords with double consonants frequently doubled and for pseudowords with an illegal position of different bigrams than for pseudowords with a legal position of these. We should note that the frontal P450 component (P300 analog) during the orthographic recognition task was only evident in the group with a high level of orthographic knowledge in Gonzalez-Garrido’s (2014) study; in our experiment, the frontal P300 effect was observed only for high-frequency words. It is considered that reading skills and word frequency affect the way we process verbal information. According to the dual-route model of reading, both direct and indirect routes can serve for semantic access (Coltheart et al., 1993; Coltheart, 2000). Through the direct route, sublexical orthographic information makes direct contact with whole-word orthographic representations. This then provide access to whole-word phonology on the one hand and higher-level semantic information on the other, unlike the direct route through an indirect, non-lexical way in which sublexical orthographic information is first transformed into a sublexical phonological code (Grainger and Ziegler, 2011). The direct way is used to recognize frequently occurring words; the meanings of more frequent words are activated by the direct route before the lower phonological route has finished processing the word (Coltheart, 2000). We assume that the P300 effect reflects mainly orthographic processing of high-frequency misspelled and correctly spelled words by the direct route and categorization processes based on orthographic properties. Taylor (1993) also noted the relation of this wave with orthography: the P300 wave was observed in the orthographic and phonological tasks, but the P300 amplitude was lower in the phonological task. Moreover, differences in P3a amplitude, a subcomponent of the P300, are observable with familiar lexical stimuli when they are orthographically unexpected (Savill and Thierry, 2011), such as misspelled words in our experiment. Interestingly, we found the P300 effect in the right anterior region, which is partially consistent with the results of Carreiras et al. (2007), who showed that the contribution of the left and right frontal areas during word and pseudoword processing are modulated by task. They found that during lexical decision task, pseudoword rejection process increases activation in the right inferior frontal cortex, and during pseudoword reading aloud, phonologic retrieval increases activation in the left precentral cortex. It is likely that during silent reading of words with and without errors, the subjects decide on the correct spelling.

We did not observe the P300 effect for low-frequency words. However, we found that a more negative response corresponding to the N400 wave was for misspelled words than for correctly spelled words and only for low-frequency words in the left posterior and right posterior regions. Previous studies have reported the association of N400 with phonological processing (Barnea and Breznitz, 1998; Zhang et al., 2009; Wang et al., 2021). Barnea and Breznitz (1998) found longer latencies and higher amplitudes of N400 during the phonological task (rhyme judgment) compared to the orthographic task (orthographic similarity/dissimilarity judgment). In most studies using pseudohomophones, phonological activation in visual word recognition and pseudohomophone effect have been associated with the N400 component, but not P300 (Kramer and Donchin, 1987; Bentin et al., 1999; Proverbio et al., 2004; Vissers et al., 2006; Briesemeister et al., 2009; González-Garrido et al., 2015; Costello et al., 2021). N400 has a larger amplitude for pseudohomophones than for words (Briesemeister et al., 2009; Hasko et al., 2013; González-Garrido et al., 2015). Misspelled words are similar to pseudohomophone stimuli; they are visually similar to correctly spelled words and are phonologically identical. Phonological plausibility causes a conflict that impedes spelling recognition, and its resolution requires repeated access to the memory, where the visual representation of the word is stored. For words presented in isolation, the N400 wave is associated with lexical–semantic processing, and the modulation of its amplitude reflects processing costs during the retrieval of properties related to a word form stored in memory (Kutas and Federmeier, 2011). The conflict caused by phonological similarity is probably more pronounced for low-frequency words and is associated with the modulation of the N400 component for misspelled words.

Phonological processing is considered slower than orthographic processing (Barnea and Breznitz, 1998). However, we showed both the N400 and P300 effects for words of different frequencies in the same time window, which seems to reflect the temporal overlap between phonological processes for low-frequency words and categorization processes based on orthographic properties for high-frequency words. Furthermore, the reaction times for high-frequency misspelled words and low-frequency misspelled words in the behavioral task did not differ. N400 can also reflect categorization processes, and some authors emphasize that, given the possible confusion of effects, it is necessary to draw careful conclusions about the processes under study (Bermúdez-Margaretto et al., 2019). On the other hand, evidence suggests that the P300 and N400 components are primarily independent and reflect two separate but interacting processes (Alday and Kretzschmar, 2019). Harm and Seidenberg (2004) proposed a cooperative division of labor between phonological and orthographic pathways to meaning activation, and word frequency may alter the relative contribution of the two routes. Some authors point out that even skilled adult readers activate the meanings of high-frequency words using phonology (Jared and O’Donnell, 2017). Thus, during spelling recognition in high-frequency and low-frequency words, two independent pathways associated with different ERP components can be active simultaneously: one reflects mainly orthographic processing and the other represents phonological processing.

It is worth noting that the process of recognizing spelling errors in our study was similar to pseudohomophone recognition. However, we got a smaller effect size than in the experiments using pseudohomophones, which were not orthographically similar to the base words (for example, stimuli “phog” and “fog” in Newman and Connolly’s study (2004); stimuli “saal” (room) and “sahl” in Braun’s study (2009)). However, the effect size was also medium in a study by González-Garrido et al. (2015), who used spelling errors as pseudohomophones. Overall, recognizing spelling errors is relatively complicated compared to recognizing pseudohomophones; nevertheless, even during passive reading, an adult native speaker can distinguish correctly spelled words from misspelled words, regardless of frequency. In addition, unlike studies in which the pseudohomophone effect was observed only for low-frequency words (Pexman et al., 2001; Braun et al., 2009), the spelling effect was observed for both high-frequency and low-frequency words and was reflected in different ERP components.

In the 350–700-ms time window, low-frequency correctly spelled words elicited a more positive effect than high-frequency correctly spelled words in left and right anterior regions. Generally, lower amplitudes for words with higher frequencies have been reported, and these results are well described (Osterhout et al., 1997; Sereno et al., 1998, 2003; Assadollahi and Pulvermüller, 2003; Hauk and Pulvermüller, 2004; Hauk et al., 2006). Nevertheless, there is little data on the relationship between the P300 component and word frequency. Polich and Donchin (1988) found larger P3 amplitudes for common words than for rare words during lexical decision tasks. However, their data could not be confirmed by Hauk and Pulvermüller’s (2004) dataset or by our results. Hauk and Pulvermüller (2004) noted differences in stimulus selection, task, data recording, and analysis as possible reasons for conflicting results. We reason Polich and Donchin’s finding (Polich and Donchin, 1988) was due to using relatively common words defined as occurring at least 30 times per million printed words and relatively uncommon words that appeared at most once per million printed words. Thus, they did not use very high-frequency words (in our experiment, every high-frequency word appears > 100 times per million) but used very low-frequency words (in our study, every low-frequency word in the ERP task appears < 3 times per million and was familiar to all participants) that could be unknown to the subjects, which may explain the differences in the results obtained. As for the timing of the word frequency effect, we expected it earlier than 350–700 ms because we used stimuli similar in length to words used in studies in which the frequency effect was observed in the time range of the P200 component (150–250 ms, e.g., Osterhout et al., 1997; Assadollahi and Pulvermüller, 2003; Dambacher et al., 2006). The frequency effect is a marker of lexical access; higher frequency words elicit a smaller amplitude than words of lower frequency, suggesting that semantic access is easier for more frequently encountered words (Van Petten and Kutas, 1990; Barber et al., 2004; Vergara-Martínez and Swaab, 2012). However, specific features of the task can affect the occurrence time of the frequency effect (Strijkers et al., 2015). Perhaps, spelling errors in words complicate lexical access; therefore, there is no difference between high-frequency and low-frequency words in the early stages. Our data also suggest that semantics access occurs in a later time window of 350–700 ms, that is, in the same window as when the processing of words with errors differs depending on the word frequency. These findings are consistent with a previous study (Wang et al., 2021), which found semantic activation in an earlier time window than we did but made similar conclusions: semantic activation of both high-frequency and low-frequency words occurs no later than phonological activation.

In the present study, we investigated the time course for recognizing the most frequent orthographic errors in Russian (error in an unstressed vowel at the root) and the effect of word frequency on this process. The findings presented here are the first of their kind to assess the time course of spelling recognition during passive reading. Based on our data and data from previous studies, we concluded that two independent pathways can be active simultaneously during spelling recognition: one mainly reflects orthographic processing for high-frequency words and the other represents phonological processing for low-frequency words. Moreover, these pathways are associated with different ERP components. Therefore, we observed the spelling effect in the same time window for both P300 and N400, which may reflect a temporal overlap mainly between categorization processes based on orthographic properties for high-frequency words and phonological processes for low-frequency words. Further research is needed on the process of recognizing fundamental errors in other languages using other paradigms (for example, the classical lexical decision task), as our results with passive reading are in many ways similar to the effects of other studies using this paradigm. In addition, it is necessary to perform source localization of ERP generators. The limitation of the current study is that we used very few electrodes, precluding any possibility of source analysis.

## Supporting information

Stimulus list

## 5 Author Contributions

EL and OM developed the general idea for the study. EL collected the data, performed the analysis, and wrote the paper. OM oversaw all stages of data analysis and edited the article.

## 8 Acknowledgments

The authors would like to thank Anna Rebreikina for her help in data collection.

