## Supplementary material for "Frequency Effects on Spelling Error Recognition: An ERP Study": Stimulus list

### *Appendix*

Table 1. Stimulus list with relevant characteristics. Behavioral task.

Note: Type – type of the stimulus; CF – correct form for HE and LE words; GN – grammatical number; Fr – frequency, ipm – instances per million words; L – length; NS – number of syllables; IPA – The International Phonetic Alphabet; EP – error position for HE and LE words

| Word | Type | CF | GN | Fr. ipm | L | NS | IPA | EP | Word translation |
|---|---|---|---|---|---|---|---|---|---|
| весна | HC | | singular | 91.3 | 5 | 2 | [vʲɪˈsna] | | spring (season) |
| глаза | HC | | plural | 857 | 5 | 2 | [ɡɫɐˈza] | | eyes |
| борьба | HC | | singular | 190.5 | 6 | 2 | [bɐrʲˈba] | | struggle, fight, combat |
| грибы | HC | | plural | 38 | 5 | 2 | [ɡrʲɪˈbɨ] | | mushrooms |
| доска | HC | | singular | 67 | 5 | 2 | [dɐˈska] | | board, plank |
| ведро | HC | | singular | 34 | 5 | 2 | [vʲɪˈdro] | | bucket, pail |
| волосы | HC | | plural | 141 | 6 | 3 | [ˈvoɫəsɨ] | | hair |
| дерево | HC | | singular | 171 | 6 | 3 | [ˈdʲerʲɪvə] | | tree, wood |
| врачи | HC | | plural | 173 | 5 | 2 | [vrɐˈt͡ɕi] | | doctors |
| плечо | HC | | singular | 236.4 | 5 | 2 | [plʲɪˈt͡ɕɵ] | | shoulder |
| стена | HC | | singular | 261 | 5 | 2 | [sʲtʲɪˈna] | | wall |
| семья | HC | | singular | 276 | 5 | 2 | [sʲɪˈmʲja] | | family |
| листы | HC | | plural | 127.6 | 5 | 2 | [lʲɪˈstɨ] | | sheets (of paper or similar) |
| места | HC | | plural | 926 | 5 | 2 | [ˈmʲestə] | | places, seats |
| плоды | HC | | plural | 47.5 | 5 | 2 | [pɫɐˈdɨ] | | fruits |
| рукава | HC | | plural | 42.9 | 6 | 3 | [rʊkɐˈva] | | sleeves |
| дворец | HC | | singular | 60 | 6 | 2 | [dvɐˈrʲet͡s] | | palace |
| добро | HC | | singular | 59 | 5 | 2 | [dɐˈbro] | | good, property |
| спина | HC | | singular | 183.1 | 5 | 2 | [spʲɪˈna] | | back |
| старик | HC | | singular | 151 | 6 | 2 | [stɐˈrʲik] | | old man |
| стекло | HC | | singular | 102.8 | 6 | 2 | [sʲtʲɪˈkɫo] | | glass |
| страна | HC | | singular | 725.7 | 6 | 2 | [strɐˈna] | | country |
| толпа | HC | | singular | 96.7 | 5 | 2 | [tɐɫˈpa] | | crowd, throng |
| ученик | HC | | singular | 80.4 | 6 | 3 | [ʊt͡ɕɪˈnʲik] | | schoolboy, pupil |
| хвосты | HC | | plural | 55.4 | 6 | 2 | [xvɐˈstɨ] | | tails |
| блины | LC | | plural | 12.8 | 5 | 2 | [blʲɪˈnɨ] | | pancakes, thin batter cakes |
| боксёр | LC | | singular | 5.3 | 6 | 2 | [bɐˈksʲɵr] | | boxer |
| винты | LC | | plural | 9.4 | 5 | 2 | [vʲɪnˈtɨ] | | screws |
| глоток | LC | | singular | 15.8 | 6 | 2 | [ɡɫɐˈtok] | | swallow, draught, gulp |
| блоха | LC | | singular | 4.3 | 5 | 2 | [bɫɐˈxa] | | flea |
| дикарь | LC | | singular | 7.4 | 6 | 2 | [dʲɪˈkarʲ] | | savage, unsociable person |
| бревно | LC | | singular | 22.5 | 6 | 2 | [brʲɪˈvno] | | log, beam (trunk of dead tree, cleared of branches) |
| дрова | LC | | plural | 25.8 | 5 | 2 | [drɐˈva] | | firewood |
| желток | LC | | singular | 3.3 | 6 | 2 | [ʐɨɫˈtok] | | yolk, vitellus |

| | | | | | | | | | |
|---|---|---|---|---|---|---|---|---|---|
| зелень | LC | | singular | 24.6 | 6 | 2 | [ˈzʲelʲɪnʲ] | | verdure (greenness, vegetation) |
| каток | LC | | singular | 7.8 | 5 | 2 | [kɐˈtok] | | skating rink, ice rink |
| крепёж | LC | | singular | 0.7 | 6 | 2 | [krʲɪˈpʲɵʂ] | | fastening, strengthening |
| кроты | LC | | plural | 6 | 5 | 2 | [krɐˈtɨ] | | moles (burrowing insectivore) |
| лебедь | LC | | singular | 14.9 | 6 | 2 | [ˈlʲebʲɪtʲ] | | swan |
| пастух | LC | | singular | 9.6 | 6 | 2 | [pɐˈstux] | | shepherd |
| пенёк | LC | | singular | 9.3 | 5 | 2 | [pʲɪˈnʲɵk] | | small stump, stub |
| вражда | LC | | singular | 7 | 6 | 2 | [vrɐˈʐda] | | enmity, hostility, animosity |
| пчела | LC | | singular | 10.3 | 5 | 2 | [pt͡ɕɪˈɫa] | | bee |
| свинья | LC | | singular | 23.1 | 6 | 2 | [svʲɪˈnʲja] | | pig, hog |
| синева | LC | | singular | 7.9 | 6 | 3 | [sʲɪnʲɪˈva] | | blue colour, blue |
| слоны | LC | | plural | 23.8 | 5 | 2 | [sɫɐˈnɨ] | | elephants |
| тропа | LC | | singular | 16.6 | 5 | 2 | [trɐˈpa] | | path, trail |
| синяк | LC | | singular | 13 | 5 | 2 | [sʲɪˈnʲak] | | bruise (medical: mark on the skin) |
| вална | HE | волна | singular | 95.4 | 5 | 2 | [vɐɫˈna] | 2 | wave |
| галова | HE | голова | singular | 709 | 6 | 3 | [gəɫɐˈva] | 2 | head |
| дажди | HE | дожди | plural | 83.2 | 5 | 2 | [dɐˈʐdʲi] | 2 | rains |
| двары | HE | дворы | plural | 166 | 5 | 2 | [dvɐˈrɨ] | 3 | courtyards |
| длена | HE | длина | singular | 67.7 | 5 | 2 | [dlʲɪˈna] | 3 | length |
| званок | HE | звонок | singular | 76.5 | 6 | 2 | [zvɐˈnok] | 3 | bell, ring, call |
| звизда | HE | звезда | singular | 122 | 6 | 2 | [zvʲɪˈzda] | 3 | star |
| зимля | HE | земля | singular | 494 | 5 | 2 | [zʲɪˈmlʲa] | 2 | earth, land, ground, soil |
| зирно | HE | зерно | singular | 30.3 | 5 | 2 | [zʲɪrˈno] | 2 | grain, seed |
| кальцо | HE | кольцо | singular | 59.5 | 6 | 2 | [kɐlʲˈt͡so] | 2 | ring, hoop |
| мазги | HE | мозги | plural | 84.5 | 5 | 2 | [mɐzˈgʲi] | 2 | brains |
| масты | HE | мосты | plural | 65 | 5 | 2 | [mɐˈstɨ] | 2 | bridges |
| нибеса | HE | небеса | plural | 33 | 6 | 3 | [nʲɪbʲɪˈsa] | 2 | heavens, skies |
| осинь | HE | осень | singular | 81 | 5 | 2 | [ˈosʲɪnʲ] | 3 | autumn, fall |
| песьмо | HE | письмо | singular | 304 | 6 | 2 | [pʲɪsʲˈmo] | 2 | letter |
| петно | HE | пятно | singular | 50 | 5 | 2 | [pʲɪtˈno] | 2 | spot, blot, stain |
| сасна | HE | сосна | singular | 30.1 | 5 | 2 | [sɐˈsna] | 2 | pine tree, pine wood |
| систра | HE | сестра | singular | 121.3 | 6 | 2 | [sʲɪˈstra] | 2 | sister |
| слиды | HE | следы | plural | 86.9 | 5 | 2 | [slʲɪˈdɨ] | 3 | tracks, footprints, traces |
| сталы | HE | столы | plural | 402.5 | 5 | 2 | [stɐˈɫɨ] | 3 | tables |
| трова | HE | трава | singular | 88.7 | 5 | 2 | [trɐˈva] | 3 | grass, herb, weed |
| халмы | HE | холмы | plural | 32.7 | 5 | 2 | [xɐɫˈmɨ] | 2 | hills |
| цвиток | HE | цветок | singular | 92.4 | 6 | 2 | [t͡svʲɪˈtok] | 3 | flower |
| чесло | HE | число | singular | 393.5 | 5 | 2 | [t͡ɕɪˈsɫo] | 2 | date, day, number |
| шкофы | HE | шкафы | plural | 48.4 | 5 | 2 | [ʂkɐˈfɨ] | 3 | cupboards, wardrobes, lockers |
| варона | LE | ворона | singular | 12.6 | 6 | 3 | [vɐˈronə] | 2 | crow (bird) |
| гниздо | LE | гнездо | singular | 20.8 | 6 | 2 | [gnʲɪˈzdo] | 3 | nest |
| граза | LE | гроза | singular | 15.7 | 5 | 2 | [grɐˈza] | 3 | thunder, thunderstorm, disaster, danger, menace |

| | | | | | | | | | |
|---|---|---|---|---|---|---|---|---|---|
| звирёк | LE | зверёк | singular | 5.3 | 6 | 2 | [zvʲɪˈrʲɵk] | 3 | small beast |
| лесица | LE | лисица | singular | 2.8 | 6 | 3 | [lʲɪˈsʲit͡sə] | 2 | fox, vixen (female of the fox) |
| лисник | LE | лесник | singular | 3.6 | 6 | 2 | [lʲɪˈsʲnʲik] | 2 | ranger, forest ranger, woodsman |
| сирьга | LE | серьга | singular | 5.7 | 6 | 2 | [sʲɪrʲˈga] | 2 | earring |
| маряк | LE | моряк | singular | 23.2 | 5 | 2 | [mɐˈrʲak] | 2 | seaman, sailor |
| начник | LE | ночник | singular | 1.6 | 6 | 2 | [nɐt͡ɕˈnʲik] | 2 | nightlight |
| митла | LE | метла | singular | 6.7 | 5 | 2 | [mʲɪˈtɫa] | 2 | broom, besom |
| повор | LE | повар | singular | 13.3 | 5 | 2 | [ˈpovər] | 4 | cook, chef |
| поруса | LE | паруса | plural | 15.2 | 6 | 3 | [pərʊˈsa] | 2 | sails |
| птинец | LE | птенец | singular | 4.6 | 6 | 2 | [ptʲɪˈnʲet͡s] | 3 | chick, nestling, baby bird |
| ражок | LE | рожок | singular | 5.1 | 5 | 2 | [rɐˈʐok] | 2 | ear trumpet, cone (of ice cream) |
| наздря | LE | ноздря | singular | 12.6 | 6 | 2 | [nɐzˈdrʲa] | 2 | nostril |
| зладей | LE | злодей | singular | 10.1 | 6 | 2 | [zɫɐˈdʲeɪ̯] | 3 | malefactor, evildoer, villain, miscreant, scoundrel |
| стрила | LE | стрела | singular | 20.7 | 6 | 2 | [strʲɪˈɫa] | 4 | arrow, pointer, indicator |
| угалёк | LE | уголёк | singular | 5.5 | 6 | 3 | [ʊgɐˈlʲɵk] | 3 | coal, ember (glowing piece of coal or wood) |
| чирвяк | LE | червяк | singular | 4.4 | 6 | 2 | [t͡ɕɪrˈvʲak] | 2 | worm |
| гарбун | LE | горбун | singular | 1.6 | 6 | 2 | [gɐrˈbun] | 2 | humpback (humpback person) |
| щепцы | LE | щипцы | plural | 1.8 | 5 | 2 | [ɕːɪpˈt͡sɨ] | 2 | tongs, pliers, pincers, nippers |
| бигун | LE | бегун | singular | 2.4 | 5 | 2 | [bʲɪˈgun] | 2 | runner, one who runs |
| лисок | LE | лесок | singular | 2.8 | 5 | 2 | [lʲɪˈsok] | 2 | grove, small wood |

Table 2. Stimulus list with relevant characteristics. ERP task.

| Word | Type | CF | GN | Fr, ipm | L | NS | IPA | EP | Word translation |
|---|---|---|---|---|---|---|---|---|---|
| война | HC | | singular | 425.9 | 5 | 2 | [vɐɪ̯ˈna] | | war |
| волосы | HC | | plural | 141 | 6 | 3 | [ˈvołəsɨ] | | hair |
| врачи | HC | | plural | 173 | 5 | 2 | [vrɐˈt͡ɕi] | | doctors |
| генерал | HC | | singular | 140 | 7 | 3 | [gʲɪnʲɪˈrał] | | general |
| глава | HC | | singular | 240.2 | 5 | 2 | [głɐˈva] | | head, top, chapter |
| желание | HC | | singular | 143.6 | 7 | 4 | [zɨ̨ˈłanʲɪɪ̯ə] | | wish, desire |
| женщина | HC | | singular | 533.3 | 7 | 3 | [ˈzɛ̨nʲɕːɪnə] | | woman |
| капитан | HC | | singular | 119 | 7 | 3 | [kəpʲɪˈtan] | | captain, commander |
| команда | HC | | singular | 174.2 | 7 | 3 | [kɐˈmandə] | | command, team |
| листы | HC | | plural | 127.6 | 5 | 2 | [lʲɪˈstɨ] | | sheets, leaves of paper or similar |
| машина | HC | | singular | 490.4 | 6 | 3 | [mɐˈʂ̨ɨnə] | | car, motor vehicle, machine |
| места | HC | | plural | 926 | 5 | 2 | [mʲɪˈsta] | | places, sites |
| министр | HC | | singular | 154.1 | 7 | 2 | [mʲɪˈnʲistr] | | minister |
| момент | HC | | singular | 306.8 | 6 | 2 | [mɐˈmʲent] | | moment |
| область | HC | | singular | 400.2 | 7 | 2 | [ˈobłəsʲtʲ] | | region, province, domain |
| офицер | HC | | singular | 118.7 | 6 | 3 | [ɐfʲɪˈt͡sɛr] | | officer |
| очередь | HC | | singular | 212.5 | 7 | 3 | [ˈot͡ɕɪrʲɪtʲ] | | queue, line, turn, order |
| период | HC | | singular | 204.3 | 6 | 3 | [pʲɪˈrʲiət] | | period, epoch, age |
| письмо | HC | | singular | 304.3 | 6 | 2 | [pʲɪsʲˈmo] | | letter |
| пример | HC | | singular | 201.2 | 6 | 2 | [prʲɪˈmʲer] | | example, instance |
| природа | HC | | singular | 169.9 | 7 | 3 | [prʲɪˈrodə] | | nature |
| радость | HC | | singular | 137.2 | 7 | 2 | [ˈradəsʲtʲ] | | joy, delight |
| ребята | HC | | plural | 148.9 | 6 | 3 | [rʲɪˈbʲatə] | | young men, boys |
| рисунок | HC | | singular | 179.2 | 7 | 3 | [rʲɪˈsunək] | | drawing, pattern |
| система | HC | | singular | 617.8 | 7 | 3 | [sʲɪˈsʲtʲemə] | | system |
| собака | HC | | singular | 132.2 | 6 | 3 | [sɐˈbakə] | | dog |
| солдат | HC | | singular | 142.2 | 6 | 2 | [sɐłˈdat] | | soldier |
| состав | HC | | singular | 209.8 | 6 | 2 | [sɐˈstaf] | | composition, structure, membership |
| спина | HC | | singular | 183.1 | 5 | 2 | [spʲɪˈna] | | back |
| старик | HC | | singular | 151 | 6 | 2 | [stɐˈrʲik] | | old man |
| статья | HC | | singular | 395 | 6 | 2 | [stɐˈtʲja] | | article, item, entry, matter, business |

| | | | | | | | |
|---|---|---|---|---|---|---|---|
| степень | HC | singular | 155 | 7 | 2 | [ˈsʲtʲepʲɪnʲ] | degree, extent, power |
| сторона | HC | singular | 768.3 | 7 | 3 | [stərɐˈna] | side |
| течение | HC | singular | 179.2 | 7 | 4 | [tʲɪˈt͡ɕenʲɪj̯ə] | current, flow, stream |
| товарищ | HC | singular | 230.6 | 7 | 3 | [tɐˈvarʲɪɕː] | comrade, friend, mate |
| элемент | HC | singular | 124.4 | 7 | 3 | [ɛlʲɪˈmʲent] | element |
| депутат | HC | singular | 108.7 | 7 | 3 | [dʲɪpʊˈtat] | deputy, delegate |
| бобёр | LC | singular | 2.6 | 5 | 2 | [bɐˈbʲɵr] | beaver |
| гармонь | LC | singular | 2.6 | 7 | 2 | [ɡɐrˈmonʲ] | garmon, accordion |
| кафель | LC | singular | 2.7 | 6 | 2 | [ˈkafʲɪlʲ] | tile |
| клубень | LC | singular | 2.7 | 7 | 2 | [ˈkɫubʲɪnʲ] | tuber |
| кочерга | LC | singular | 2.8 | 7 | 3 | [kət͡ɕɪrˈɡa] | poker |
| лазейка | LC | singular | 2.7 | 7 | 3 | [ɫɐˈzʲej̯kə] | narrow hole, loophole |
| летун | LC | singular | 0.9 | 5 | 2 | [lʲɪˈtun] | flyer |
| лилипут | LC | singular | 2.7 | 7 | 3 | [lʲɪlʲɪˈput] | Lilliputian, dwarf |
| литера | LC | singular | 2.7 | 6 | 3 | [ˈlʲitʲɪrə] | letter, character |
| лифтёр | LC | singular | 1 | 6 | 2 | [lʲɪˈftʲɵr] | elevator boy, lift man |
| ловец | LC | singular | 2.4 | 5 | 2 | [ɫɐˈvʲet͡s] | fisherman, hunter |
| логово | LC | singular | 2.8 | 6 | 3 | [ˈɫoɡəvə] | den, lair |
| лодочка | LC | singular | 2.6 | 7 | 3 | [ˈɫodət͡ɕkə] | small boat |
| лосиха | LC | singular | 0.6 | 6 | 3 | [ɫɐˈsʲixə] | female moose |
| лужица | LC | singular | 2.6 | 6 | 3 | [ˈɫuʐɨt͡sə] | small puddle, pool |
| манжета | LC | singular | 2.7 | 7 | 3 | [mɐnˈʐɛtə] | cuff, wristband |
| метраж | LC | singular | 1.1 | 6 | 2 | [mʲɪˈtraʂ] | footage, length in meters, area in square meters |
| мирок | LC | singular | 2 | 5 | 2 | [mʲɪˈrok] | small world, microcosm |
| немота | LC | singular | 2.7 | 6 | 3 | [nʲɪmɐˈta] | muteness, dumbness |
| новатор | LC | singular | 2.8 | 7 | 3 | [nɐˈvatər] | innovator |
| обивка | LC | singular | 2.9 | 6 | 3 | [ɐˈbʲifkə] | upholstery |
| объятье | LC | singular | 2.6 | 7 | 3 | [ɐˈbjætʲɪj̯ə] | embrace (hug) |
| огласка | LC | singular | 2.8 | 7 | 3 | [ɐˈɡɫaskə] | publicity |
| огрызок | LC | singular | 2.6 | 7 | 3 | [ɐˈɡrɨzək] | stub, stump |
| озерцо | LC | singular | 0.8 | 6 | 3 | [ɐzʲɪrˈt͡so] | small lake |
| пальба | LC | singular | 2.9 | 6 | 2 | [pɐlʲˈba] | shooting |
| перелив | LC | singular | 2.6 | 7 | 3 | [pʲɪrʲɪˈlʲif] | overflow |
| перина | LC | singular | 2.4 | 6 | 3 | [pʲɪˈrʲinə] | a feather bed |
| пилюля | LC | singular | 2.7 | 6 | 3 | [pʲɪˈlʲʉlʲə] | pill, pilule |

| | | | | | | | | | |
|---|---|---|---|---|---|---|---|---|---|
| писарь | LC | | singular | 2.9 | 6 | 2 | [ˈpʲisərʲ] | | scribe, scrivener |
| разнос | LC | | singular | 2.8 | 6 | 2 | [rɐˈznos] | | carrying, delivery |
| раскат | LC | | singular | 2.6 | 6 | 2 | [rɐˈskat] | | reverberation, roll (a heavy, reverberatory sound) |
| синица | LC | | singular | 2.9 | 6 | 3 | [sʲɪˈnʲit͡sə] | | titmouse, tomtit |
| слякоть | LC | | singular | 2.7 | 7 | 2 | [ˈslʲakətʲ] | | slush, mire, crud, mud |
| солярка | LC | | singular | 2.9 | 7 | 3 | [sɐˈlʲarkə] | | diesel fuel |
| сонет | LC | | singular | 2.8 | 5 | 2 | [sɐˈnʲet] | | sonnet |
| тирада | LC | | singular | 2.8 | 6 | 3 | [tʲɪˈradə] | | tirade |
| трагизм | LC | | singular | 2.7 | 7 | 2 | [trɐˈɡʲizm] | | tragedy, tragic element |
| абъект | HE | объект | singular | 206.4 | 6 | 2 | [ɐˈbjekt] | 1 | object |
| барьба | HE | борьба | singular | 190.5 | 6 | 2 | [bɐrʲˈba] | 1 | struggle, fight, combat |
| вапрос | HE | вопрос | singular | 805.8 | 6 | 2 | [vɐˈpros] | 1 | question |
| влеяние | HE | влияние | singular | 114.9 | 7 | 3 | [vlʲɪˈjænʲɪɪ̯ə] | 1 | influence, effect, weight, credibility |
| выбары | HE | выборы | plural | 117.7 | 6 | 3 | [ˈvɨbərɨ] | 2 | election |
| вывад | HE | вывод | singular | 111.8 | 5 | 2 | [ˈvɨvət] | 2 | conclusion, inference, deduction |
| девачка | HE | девочка | singular | 185.1 | 7 | 3 | [ˈdʲevət͡ɕkə] | 2 | girl, female child |
| дериво | HE | дерево | singular | 171 | 6 | 3 | [ˈdʲerʲɪvə] | 2 | tree, wood |
| диревня | HE | деревня | singular | 125.1 | 7 | 3 | [dʲɪˈrʲevnʲə] | 1 | village, hamlet |
| доктар | HE | доктор | singular | 143.1 | 6 | 2 | [ˈdoktər] | 2 | doctor |
| зимля | HE | земля | singular | 494.4 | 5 | 2 | [zʲɪˈmlʲa] | 1 | earth, land, ground, soil |
| интирес | HE | интерес | singular | 260.6 | 7 | 3 | [ɪnʲtʲɪˈrʲes] | 2 | interest |
| кабенет | HE | кабинет | singular | 148.5 | 7 | 3 | [kəbʲɪˈnʲet] | 2 | office |
| карабль | HE | корабль | singular | 112.5 | 7 | 2 | [kɐˈrablʲ] | 1 | ship |
| мадель | HE | модель | singular | 153.5 | 6 | 2 | [mɐˈdɛlʲ] | 1 | model |
| пабеда | HE | победа | singular | 124 | 6 | 3 | [pɐˈbʲedə] | 1 | victory |
| панятие | HE | понятие | singular | 139.4 | 7 | 4 | [pɐˈnʲætʲɪɪ̯ə] | 1 | idea, concept, notion |
| паринь | HE | парень | singular | 140.3 | 6 | 2 | [ˈparʲɪnʲ] | 2 | fellow, lad, chap, guy, boyfriend |
| парядок | HE | порядок | singular | 307.6 | 7 | 3 | [pɐˈrʲadək] | 1 | order, sequence |
| пириод | HE | период | singular | 204.3 | 6 | 3 | [pʲɪˈrʲiət] | 1 | period, epoch |
| пличо | HE | плечо | singular | 236.4 | 5 | 2 | [plʲɪˈt͡ɕɵ] | 1 | shoulder |
| правело | HE | правило | singular | 258.8 | 7 | 3 | [ˈpravʲɪɫə] | 2 | rule, regulations, law, canon |
| прадукт | HE | продукт | singular | 136.7 | 7 | 2 | [prɐˈdukt] | 1 | product |
| пречина | HE | причина | singular | 237.2 | 7 | 3 | [prʲɪˈt͡ɕinə] | 1 | cause |

| | | | | | | | | | |
|---|---|---|---|---|---|---|---|---|---|
| придмет | HE | предмет | singular | 154.4 | 7 | 2 | [prʲɪdˈmʲet] | 1 | object, subject, topic |
| рибёнок | HE | ребёнок | singular | 658.3 | 7 | 3 | [rʲɪˈbʲɵnək] | 1 | child, kid, baby |
| ригион | HE | регион | singular | 160.3 | 6 | 3 | [rʲɪgʲɪˈon] | 1 | region |
| ришение | HE | решение | singular | 453.4 | 7 | 4 | [rʲɪˈʂɛnʲɪ̯ə] | 1 | decision, solution, answer |
| сабытие | HE | событие | singular | 206.5 | 7 | 4 | [sɐˈbɨtʲɪ̯ə] | 1 | event, occurrence, incident |
| свабода | HE | свобода | singular | 174.9 | 7 | 3 | [svɐˈbodə] | 1 | freedom, liberty |
| симья | HE | семья | singular | 276 | 5 | 2 | [sʲɪˈmʲja] | 1 | family |
| слиза | HE | слеза | singular | 114.2 | 5 | 2 | [slʲɪˈza] | 1 | tear |
| стикло | HE | стекло | singular | 102.8 | 6 | 2 | [sʲtʲɪˈkło] | 1 | glass |
| стина | HE | стена | singular | 261 | 5 | 2 | [sʲtʲɪˈna] | 1 | wall |
| тилефон | HE | телефон | singular | 167.8 | 7 | 3 | [tʲɪlʲɪˈfon] | 1 | telephone, telephone number |
| тиория | HE | теория | singular | 116.5 | 6 | 4 | [tʲɪˈorʲɪ̯ə] | 1 | theory |
| тысича | HE | тысяча | singular | 416 | 6 | 3 | [ˈtɨsʲɪt͡ɕə] | 2 | thousand |
| уравень | HE | уровень | singular | 348.5 | 7 | 3 | [ˈurəvʲɪnʲ] | 2 | level, standard, amount |
| хазяин | HE | хозяин | singular | 170.6 | 6 | 3 | [xɐˈzʲaɪn] | 1 | owner, proprietor |
| абрезок | LE | обрезок | singular | 2.7 | 7 | 3 | [ɐˈbrʲezək] | 1 | end, shred, snippet |
| абслуга | LE | обслуга | singular | 2.9 | 7 | 3 | [ɐpˈsługə] | 1 | staff, service personnel |
| аракул | LE | оракул | singular | 2.7 | 6 | 3 | [ɐˈrakʊł] | 1 | oracle |
| аткат | LE | откат | singular | 2.9 | 5 | 2 | [ɐˈtkat] | 1 | recoil, retreat |
| бахрама | LE | бахрома | singular | 2.6 | 7 | 3 | [bəxrɐˈma] | 2 | fringe (decorative border) |
| бегатня | LE | беготня | singular | 2.7 | 7 | 3 | [bʲɪgɐtʲˈnʲa] | 2 | running about, bustle |
| белетик | LE | билетик | singular | 2.6 | 7 | 3 | [bʲɪˈlʲetʲɪk] | 1 | ticket |
| београф | LE | биограф | singular | 2.6 | 7 | 3 | [bʲɪˈogrəf] | 1 | biographer |
| варонок | LE | воронок | singular | 2.6 | 7 | 3 | [vərɐˈnok] | 1 | common house martin, paddy wagon |
| гарняк | LE | горняк | singular | 2.6 | 6 | 2 | [gɐrˈnʲak] | 1 | miner |
| гомак | LE | гамак | singular | 2.9 | 5 | 2 | [gɐˈmak] | 1 | hammock |
| двежок | LE | движок | singular | 2.9 | 6 | 2 | [dvʲɪˈʐok] | 1 | engine, motor |
| деалект | LE | диалект | singular | 2.9 | 7 | 3 | [dʲɪɐˈlʲekt] | 1 | dialect |
| ельнек | LE | ельник | singular | 2.6 | 6 | 2 | [ˈjelʲnʲɪk] | 2 | fir-grove, fir forest |
| заветок | LE | завиток | singular | 2.9 | 7 | 3 | [zəvʲɪˈtok] | 2 | curl, swirl, spiral |
| зивота | LE | зевота | singular | 1.2 | 6 | 3 | [zʲɪˈvotə] | 1 | yawning |
| кодет | LE | кадет | singular | 2.9 | 5 | 2 | [kɐˈdʲet] | 1 | cadet |

| | | | | | | | | | |
|---|---|---|---|---|---|---|---|---|---|
| крохмал | LE | крахмал | singular | 2.9 | 7 | 2 | [krɐˈxmaɫ] | 1 | starch |
| матылёк | LE | мотылёк | singular | 2.7 | 7 | 3 | [mətɨˈlʲɵk] | 1 | moth, butterfly |
| мегалка | LE | мигалка | singular | 2.8 | 7 | 3 | [mʲɪˈgaɫkə] | 1 | blinkenlight, emergency vehicle lighting |
| ногота | LE | нагота | singular | 2.7 | 6 | 3 | [nəgɐˈta] | 1 | nudity, bareness, nakedness |
| пакрой | LE | покрой | singular | 2.9 | 6 | 2 | [pɐˈkroɪ̯] | 1 | cut, style (of garments) |
| пекет | LE | пикет | singular | 2.7 | 5 | 2 | [pʲɪˈkʲet] | 1 | picket |
| питерня | LE | пятерня | singular | 2.7 | 7 | 3 | [pʲɪtʲɪrˈnʲa] | 1 | all five fingers, palm with five fingers |
| прагон | LE | прогон | singular | 2.8 | 6 | 2 | [prɐˈgon] | 1 | driving of animals, architecture purlin |
| прареха | LE | прореха | singular | 2.7 | 7 | 3 | [prɐˈrʲexə] | 1 | tear, slit, hole, lapse, gap |
| рефират | LE | реферат | singular | 2.9 | 7 | 3 | [rʲɪfʲɪˈrat] | 2 | abstract, synopsis, summary |
| римарка | LE | ремарка | singular | 2.8 | 7 | 3 | [rʲɪˈmarkə] | 1 | remark, note |
| роздор | LE | раздор | singular | 2.8 | 6 | 2 | [rɐˈzdor] | 1 | discord, contention, dissension |
| розлад | LE | разлад | singular | 2.9 | 6 | 2 | [rɐˈzɫat] | 1 | discord, dissension |
| сеница | LE | синица | singular | 2.9 | 6 | 3 | [sʲɪˈnʲit͡sə] | 1 | titmouse, tomtit |
| сидмица | LE | седмица | singular | 2.6 | 7 | 3 | [sʲɪdʲˈmʲit͡sə] | 1 | week |
| симестр | LE | семестр | singular | 2.8 | 7 | 2 | [sʲɪˈmʲestr] | 1 | semester, term (half of school year) |
| скокун | LE | скакун | singular | 1.9 | 6 | 2 | [skɐˈkun] | 1 | racehorse |
| фетиль | LE | фитиль | singular | 2.7 | 6 | 2 | [fʲɪˈtʲilʲ] | 1 | wick, fuse |
| фригат | LE | фрегат | singular | 2.8 | 6 | 2 | [frʲɪˈgat] | 1 | frigate, frigatebird |
| хохалок | LE | хохолок | singular | 2.6 | 7 | 3 | [xəxɐˈɫok] | 2 | crest, topknot |
| ямачка | LE | ямочка | singular | 2.8 | 6 | 3 | [ˈjamət͡ɕkə] | 2 | small hole, pit, socket |
| ясинь | LE | ясень | singular | 2.6 | 5 | 2 | [ˈjæsʲɪnʲ] | 2 | ash |
